# 2020 BioImage Analysis Survey: Community experiences and needs for the future

**DOI:** 10.1101/2021.08.16.456498

**Authors:** Nasim Jamali, Ellen TA Dobson, Kevin W. Eliceiri, Anne E. Carpenter, Beth A. Cimini

## Abstract

In this paper, we summarize a global survey of 484 participants of the imaging community, conducted in 2020 through the NIH Center for Open BioImage Analysis (COBA). This 23-question survey covered experience with image analysis, scientific background and demographics, and views and requests from different members of the imaging community. Through open-ended questions we asked the community to provide feedback for the open-source tool developers and tool user groups. The community’s requests for tool developers include general improvement of tool documentation and easy-to-follow tutorials. Respondents encourage tool users to follow the best practices guidelines for imaging and ask their image analysis questions on the Scientific Community Image forum (forum.image.sc). We analyzed the community’s preferred method of learning, based on level of computational proficiency and work description. In general, written step-by-step and video tutorials are preferred methods of learning by the community, followed by interactive webinars and office hours with an expert. There is also enthusiasm for a centralized location online for existing educational resources. The survey results will help the community, especially developers, trainers, and organizations like COBA, decide how to structure and prioritize their efforts.

**Impact statement:** The Bioimage analysis community consists of software developers, imaging experts, and users, all with different expertise, scientific background, and computational skill levels. The NIH funded Center for Open Bioimage Analysis (COBA) was launched in 2020 to serve the cell biology community’s growing need for sophisticated open-source software and workflows for light microscopy image analysis. This paper shares the result of a COBA survey to assess the most urgent ongoing needs for software and training in the community and provide a helpful resource for software developers working in this domain. Here, we describe the state of open-source bioimage analysis, developers’ and users’ requests from the community, and our resulting view of common goals that would serve and strengthen the community to advance imaging science.

## Results

### Participants

Of the 484 survey participants, the majority were from North America (60%) and Europe (34%), followed by Asia, Australia, South America, and Africa (Supplementary Fig 1A). 43% of participants were in training (either “postdoctoral fellow” or “undergrad/graduate students”, hereafter described as “Trainees”; Figure 1A). Most participants described experience and/or training in the biological sciences, including cell/molecular biology, chemistry/biochemistry, followed by physics and developmental biology (Figure 1B), though the small proportion of participants whose role was “Image analyst” or “Other” reported computational backgrounds more often. This is consistent with a 2015 survey from the Network of European BioImage Analysts (NEUBIAS) ^1^, which is a network working toward bridging the efforts between life science, computer science, and digital image processing ^2^.

**Figure 1.**
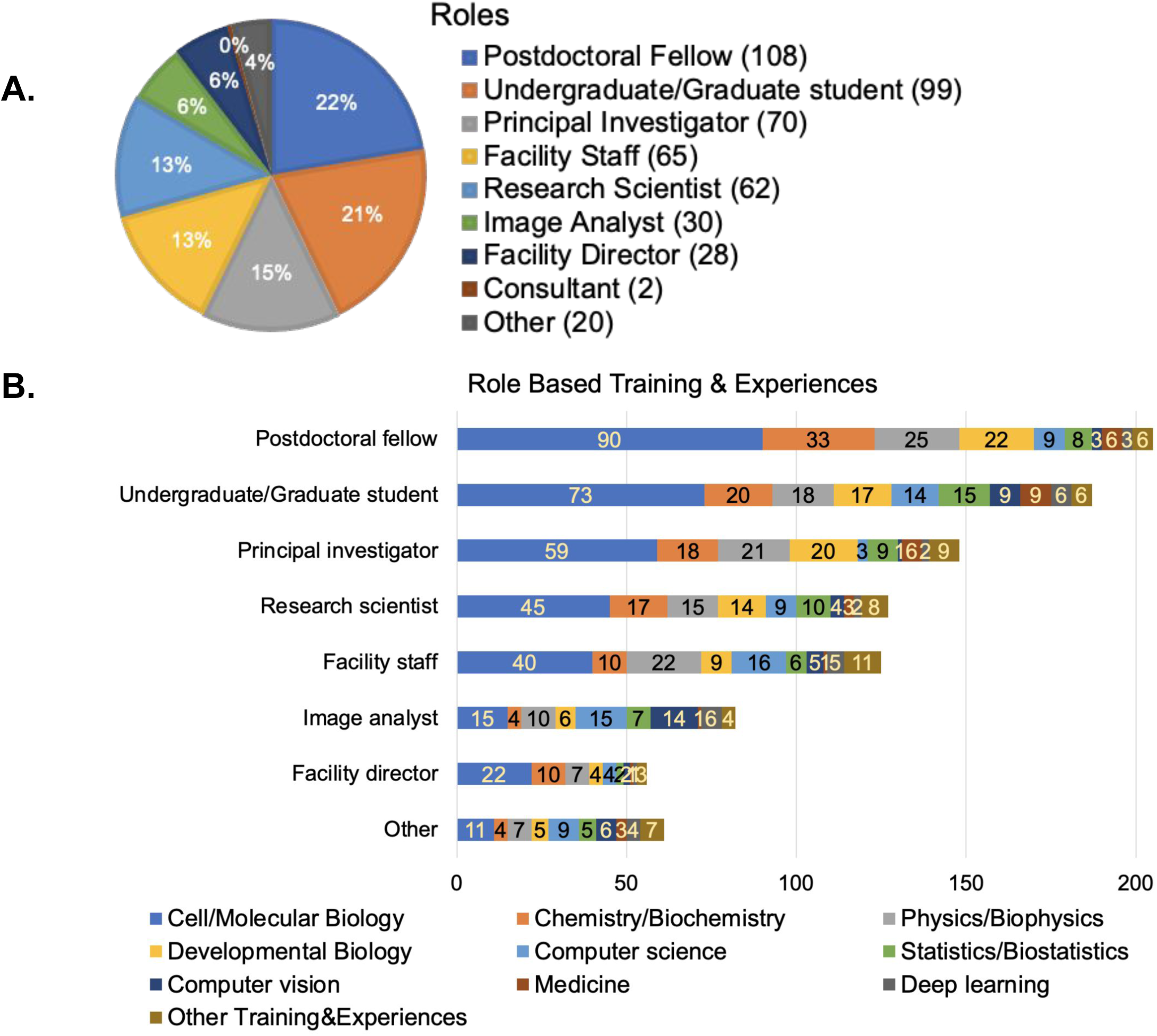
A. Responses to the multi-choice question of “Which of the following roles best describe you?”. Respondents had the option to provide additional answers. B. Responses to the checkbox question of “Which of the following do you have significant formal training in or experience with? Select all that apply”. Responses were broken down by the major categories from part A.

We next asked participants to describe their work along a linear scale of 1 to 7, with 1 representing mainly imaging (sample preparation, optimizing/deciding on imaging modalities, acquiring images, etc.) and 7 representing mainly image analysis (finding the right tools to analyze a particular experiment, optimizing the analysis, and data mining). The group of respondents with scores of 1 and 2 are hereafter described as “imaging”, the group with scores of 3-5 are described as “balanced”, and the group with scores of 6-7 as “analysts”. The majority of respondents fell into the “balanced” category (Supplementary Figure 1B).

We also asked participants about both their current level of computational skills and comfort developing new skills on a scale of 1 to 7, with 1 representing very poor and 7 representing excellent for current skills and 1 representing very uncomfortable and 7 representing very comfortable. Computational skills tracked somewhat with job roles, with the imaging, balanced, and analyst groups showing mean computational skills of 3.59, 4.17, and 5.40, respectively (Figure 2). On average, the community feels they are more comfortable than not in developing new computational skills, rating themselves a 4.73 out of 7 (Supplementary Figure 1C).

**Figure 2.**
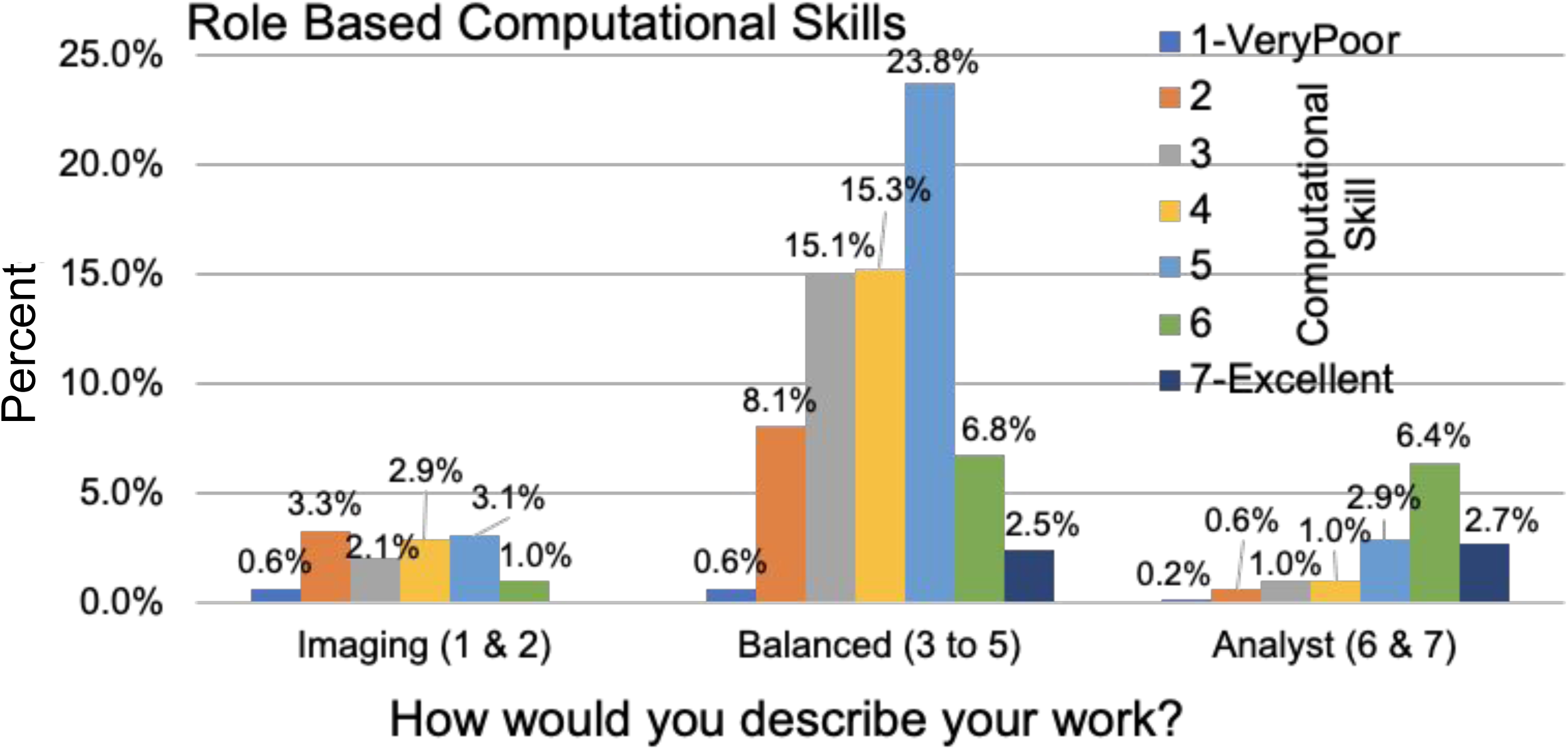
Responses to the question of “How would you rate your computational skills?” along a linear scale with 1 representing Very poor and 7 representing Excellent. Responses were cross-matched with the work duties (graphed in Supplementary Figure 1B).

Since almost half of the survey participants were from the trainee group and because trainees may have particular requests that need to be addressed, we wanted to know how our measures of job class, computational skill, and computational comfort differed for trainees vs non-trainees. Cross-matching the work duties as shown in Supplementary Figure 1B with the role description as shown in Figure 1A, showed the trainee group was relatively evenly drawn from the imaging, balanced, and analyst groups. A large number of survey respondents were from the balanced group, regardless of their trainee status (Figure 3A). While trainees were missing from the highest self-reported skill level, reported comfort in developing new computational skills increased with reported existing skill level and was comparable between trainees and non-trainees (Figure 3B).

**Figure 3.**
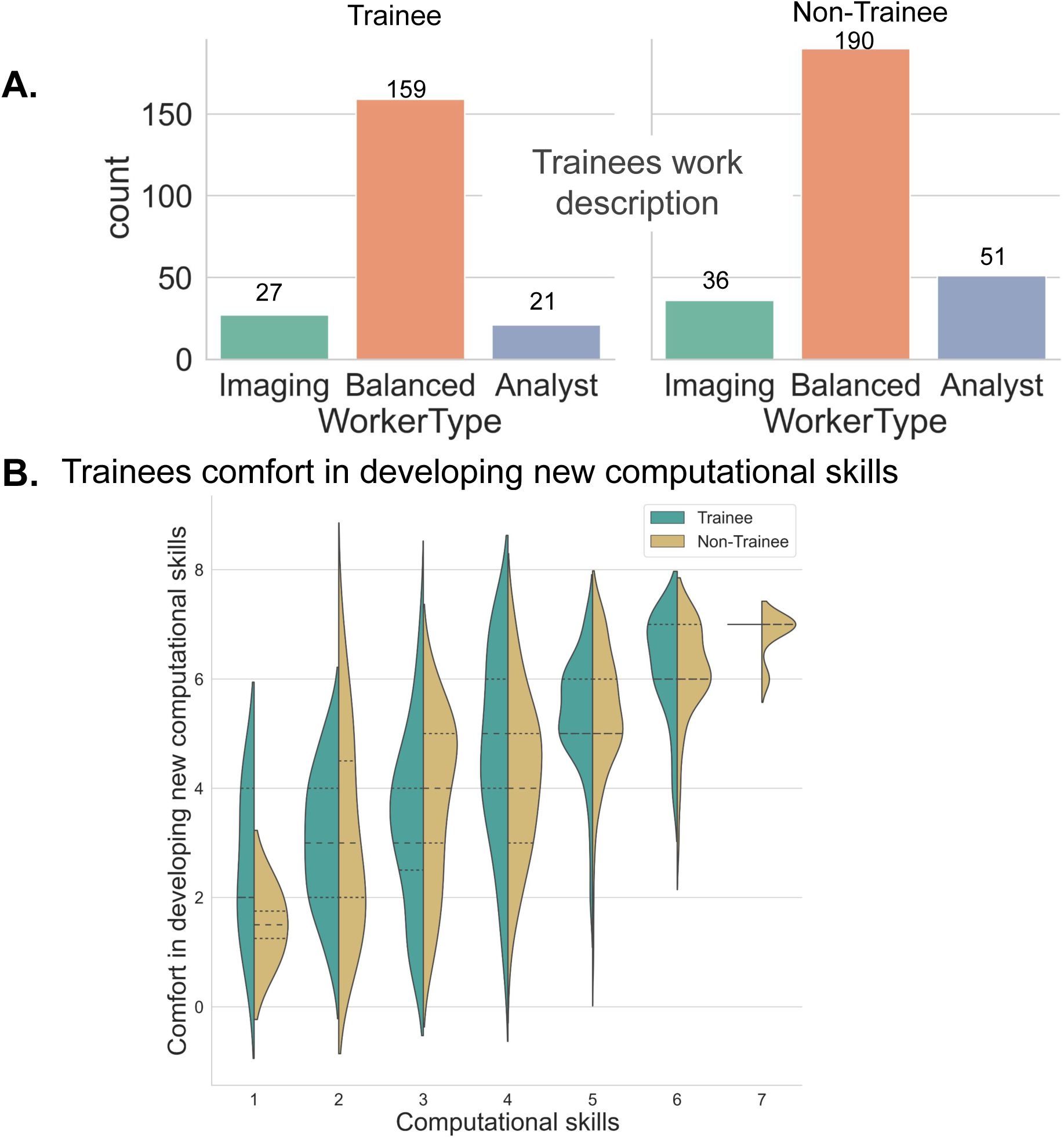
A. The trainee vs non-trainee groups’ job description were drawn by cross-match of the work duties (graphed in Supplementary Figure 1B) and role description (graphed in Figure 1A) B. Responses to the question of “How would you rate your comfort in developing new computational skills?” (graphed in Supplementary Figure 1C), cross-matched with the responses to the question of “How would you rate your computational skills?”, and separated based on the trainee group.

### Needs and commonly-used tools

#### Types of images analyzed (modality, technique, specimen, etc.)

In response to our question “*What kinds of images do you commonly want to analyze?*”, brightfield (BF/DIC/Phase) and fluorescent two-dimensional (2D) images from cells and organisms were the top most-commonly analyzed images among the survey participants (Figure 4), which were mainly captured by manual field selection as opposed to an automated system. Overall the majority of images to be analyzed were 2D, followed by 2D timelapse images, 3D volumes, and then 3D timelapse images.

**Figure 4.**
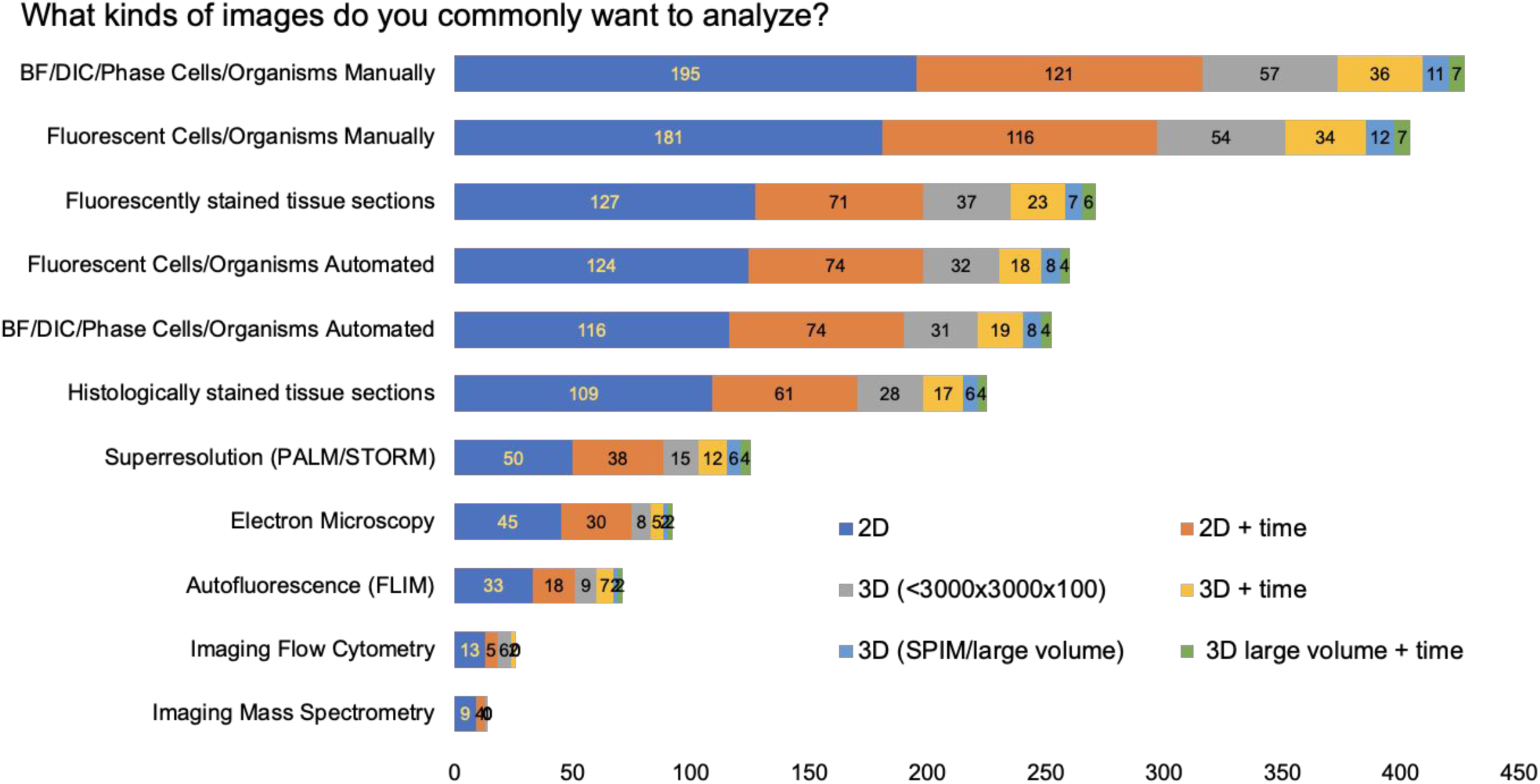
Responses to the checkbox grid question of “What kinds of images do you commonly want to analyze?”. Respondents had the option to provide additional answers.

#### Types of analysis tools used

When we asked “*What image analysis tools do you use the most?*”, the vast majority of respondents reported the category of “open source point and click software”. The question described such tools to include ImageJ ^3,4^, Fiji ^5^, CellProfiler ^6^, Icy ^7^, etc., all of which are the most commonly used tools among participants (Figure 5A). The next most common tools were “computational libraries and scripts” (such as scikit-image ^8^ and MATLAB ^9^ libraries), followed by commercial software on the user’s microscope and other commercial software. While these may be representative of the broader community, we note that most promotion of the survey was via a forum for open-source tools (forum.image.sc) ^10^ and by Twitter accounts related to open-source tools, presumably increasing their final representation in the survey.

**Figure 5.**
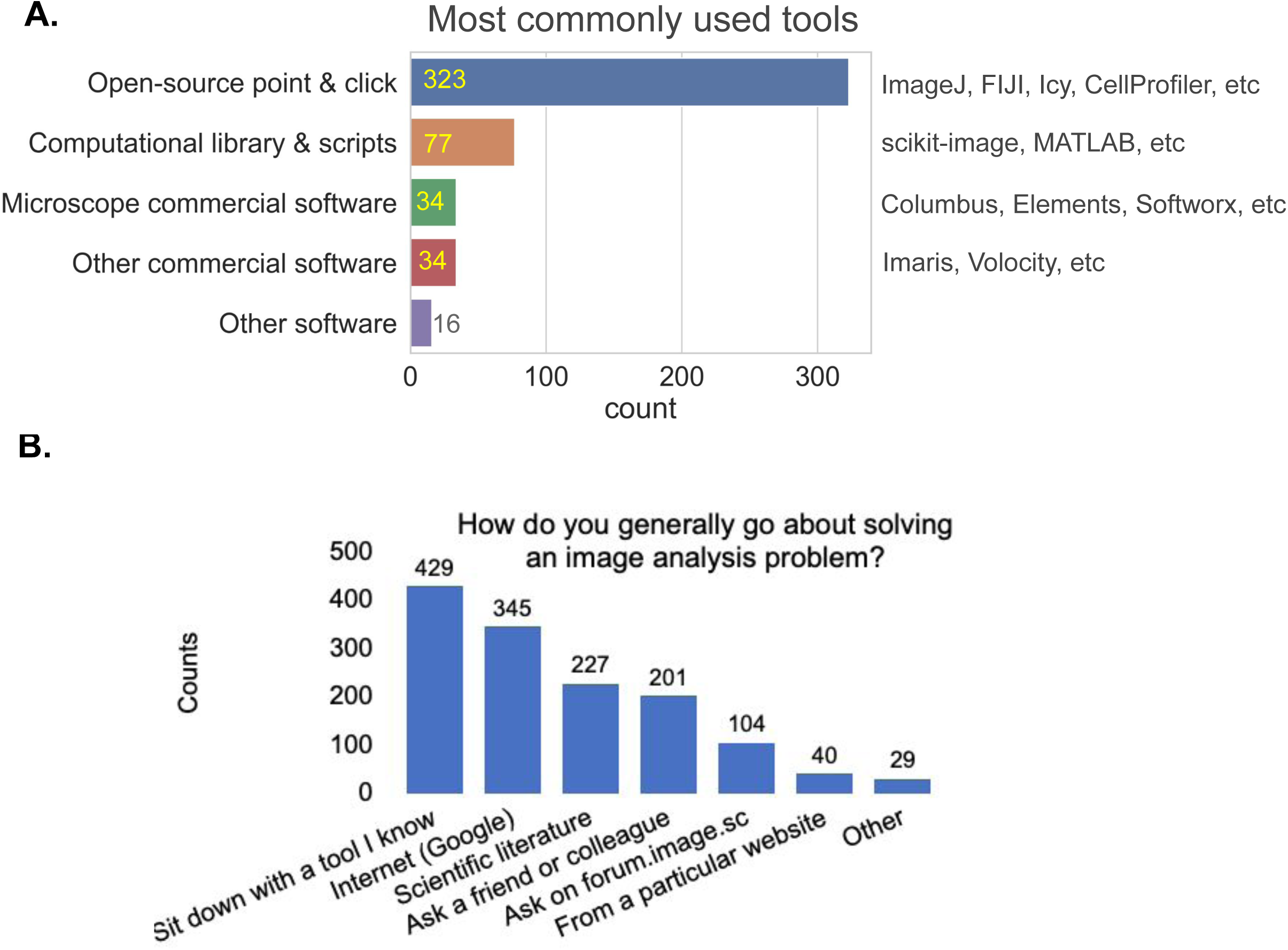
A. Responses to the multiple choice question of “What image analysis tools do you use the most?”. Respondents had the option to provide additional options. B. Responses to the checkbox question of “How do you generally go about solving an image analysis problem? Check the approach(es) you use them the most”. Respondents had the option to provide additional options.

### Approaches, image analysis problems, and tips for developers and users

#### Commonly used approaches to solve image analysis problems

We next wanted to better understand what approaches our participants take to solve their image analysis problems. The most popular approaches are: sit down with a familiar tool, search the web, look up solutions in the scientific literature, ask a friend or colleague, and/or ask on the Scientific Community Image forum (forum.image.sc) (Figure 5B).

#### Well-solved image analysis issues and those needing better solutions

To assess the community’s thoughts on the success of currently existing tools, we next asked participants via open-ended questions to describe which image analysis problems they thought were already well-solved and which they wished had better solutions. We analyzed the free-text responses by parsing them, using Python scripts as described in the methods section. Responses containing “detection” and/or “segmentation” were the most common for both questions; likely this reflects the huge diversity of biological objects our participants are trying to detect (Figure 6A-B). The answers with a much higher relative rank in the “well-solved” category were “Nuclei” and “2D” and in the “wish were better solved” category were “3D/Volume” and “Tissue/Histology’’.

**Figure 6.**
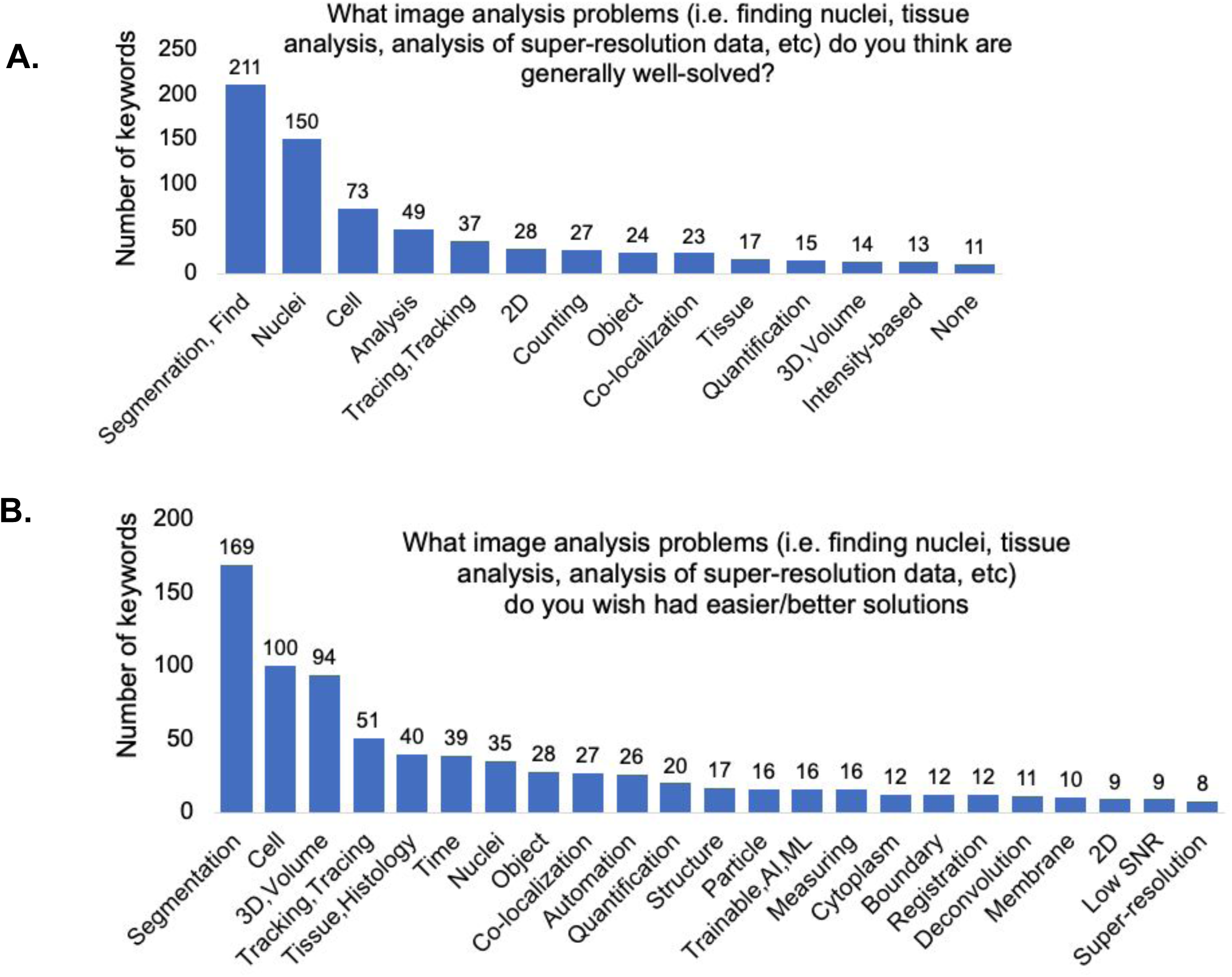
A. Keyword analysis of the open-ended question of “What image analysis problems (i.e. finding nuclei, tissue analysis, analysis of super-resolution data, etc) do you think are generally well-solved?”. B. Keyword analysis of the open-ended question of “What image analysis problems (i.e. finding nuclei, tissue analysis, analysis of super-resolution data, etc) do you wish had easier/better solutions?”.

### Tips for image analysis for users and developers

#### Suggestions for the Open-Source Tool Creators/ What creators can do

We asked the community to provide open-ended feedback for the tool developers: “*What do you think analysis tool CREATORS (such as software developers) could/should do to make image analysis better and more successful? How best could we encourage them to do it?*”. Through our analysis of keywords, we conclude that all groups encouraged developers to improve documentations and manuals, user interface, plugins and packages, as well as providing easy-to-follow workshops and tutorials (Figure 7A). They suggested that tool creators form a tight-knit community, in which they could collaborate, review software, and improve documentation. Other suggestions included prizes for best tool instructions and frequent reminders for users to cite analysis tools in publications.

**Figure 7.**
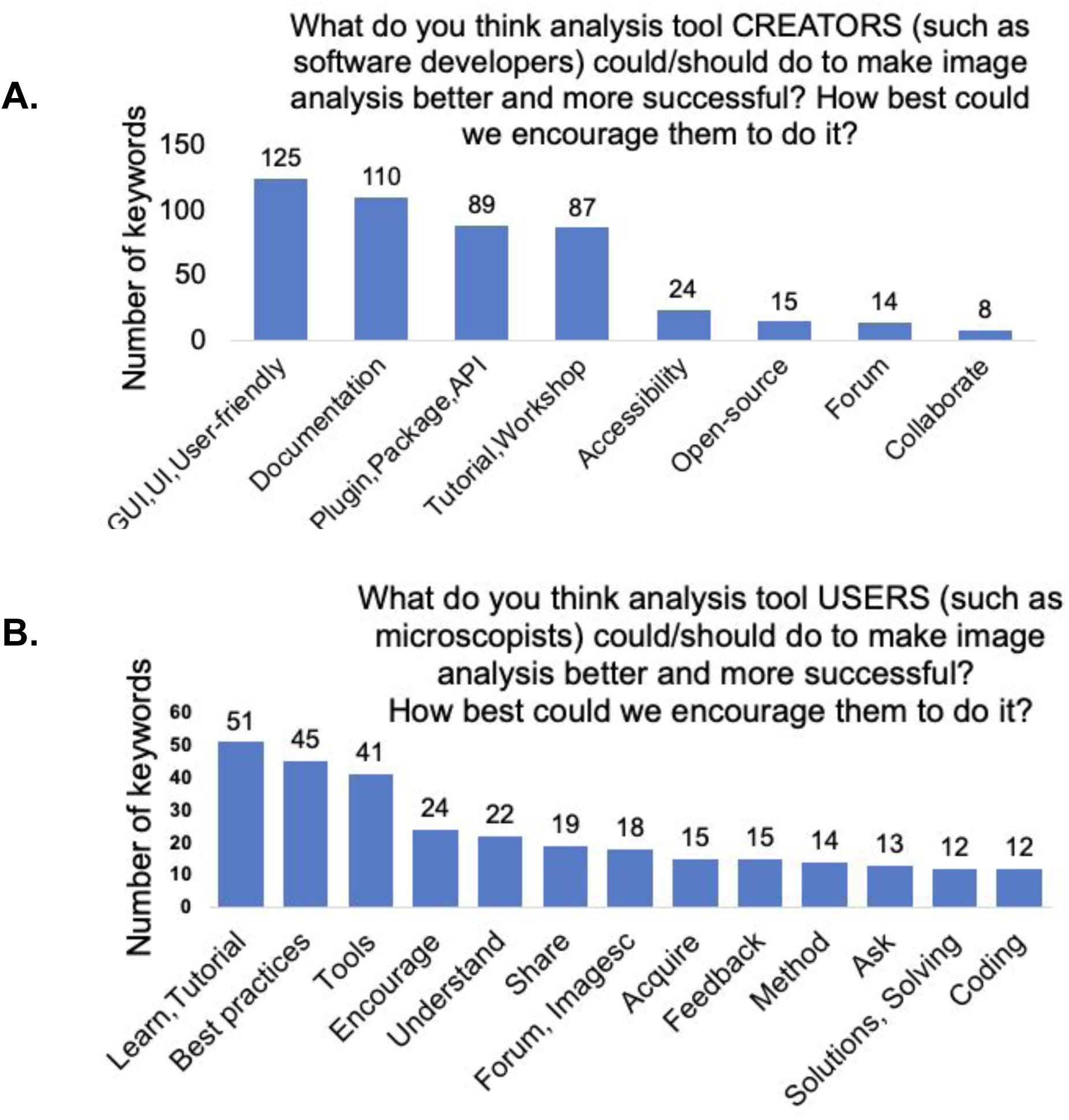
A. Keyword analysis of the open-ended question of “What do you think analysis tool CREATORS (such as software developers) could/should do to make image analysis better and more successful? How best could we encourage them to do it?”. B. Keyword analysis of the open-ended question of “What do you think analysis tool USERS (such as microscopists) could/should do to make image analysis better and more successful? How best could we encourage them to do it?”.

As we expected, users who self-reported spending more time on imaging than on analysis have different perspectives on the greatest needs. We therefore binned the answers into the “imaging”, “balanced”, and “analyst” categories. The “imaging” group’s most common suggestions were for more-user friendly tools, better documentation, and more video tutorials. The “balanced” group emphasized a need for clear documentation with detailed explanations of the parameters, as well as asking for tools that perform better in real-world conditions when ability to improve sample acquisition is limited. The “analysts” also highlighted a need for better documentation, an ecosystem with fewer and more multifunctional tools over a plethora of single-use scripts, and more interoperability between the current open source tools; they also encouraged developers to communicate with users early and often.

#### Suggestions for the Open-Source Tool Users

We also asked, “*What do you think analysis tool USERS (such as microscopists) could/should do to make image analysis better and more successful? How best could we encourage them to do it?*”, allowing the community to provide feedback for the tool users. Overall, as shown in Figure 7B, the community asked users to seek out existing tutorials and workshops and learn the tools and basics of image analysis, follow the best practices guidelines for imaging, and ask their image analysis questions on the image.sc forum. By using a single, central forum for image analysis questions, all members could benefit from every user’s questions, feedback, and suggestions, which makes it easier for tool creators to provide answers. Respondents also encouraged users to start image analysis when performing pilot imaging and condition optimization experiments, rather than waiting until the completion of larger-scale experiments. Participants also suggested that principal investigators encourage their trainees to take image analysis courses and try new image analysis tools.

Breaking down the answers by role, we found that the “imaging” group asked the users to pay attention to sample quality, to have an understanding of why particular image analysis methods are applied in certain cases, and to reach out to experts and provide detailed feedback to tool developers with examples. The “balanced” group responded that users should provide analysis workflow details in publications, keep settings consistent during acquisition and analysis, and they encouraged tool users to think about the analysis before and during image acquisition, testing their approach on early data rather than leaving all the analysis for the end. They also emphasized providing feedback for the developers, exploring new tools, and posting questions on the image.sc forum. “Analysts” particularly asked users to take advantage of available tutorials and workshops about the basics of image analysis and open source tools. They also encouraged tool users to clarify their image analysis needs to the developers, provide feedback, request new features, and report bugs, including detailed information and sample images in their bug reports.

### Learning materials and their components

#### Interest in learning new topics

We next asked users to rate their interest in learning more about each of several topics. Participants showed the highest level of interest in learning about image analysis practices specifically related to their own field (Figure 8A). The remaining four areas to be rated, image analysis theories, general image analysis practices, learning to use different software tools, and deep learning approaches for image analysis, showed a similar pattern to each other with a plurality of users stating they were “very interested” followed by “moderately”, “a little”, and “not at all” interested. These results emphasize that the community shows an appetite for image analysis educational materials.

**Figure 8.**
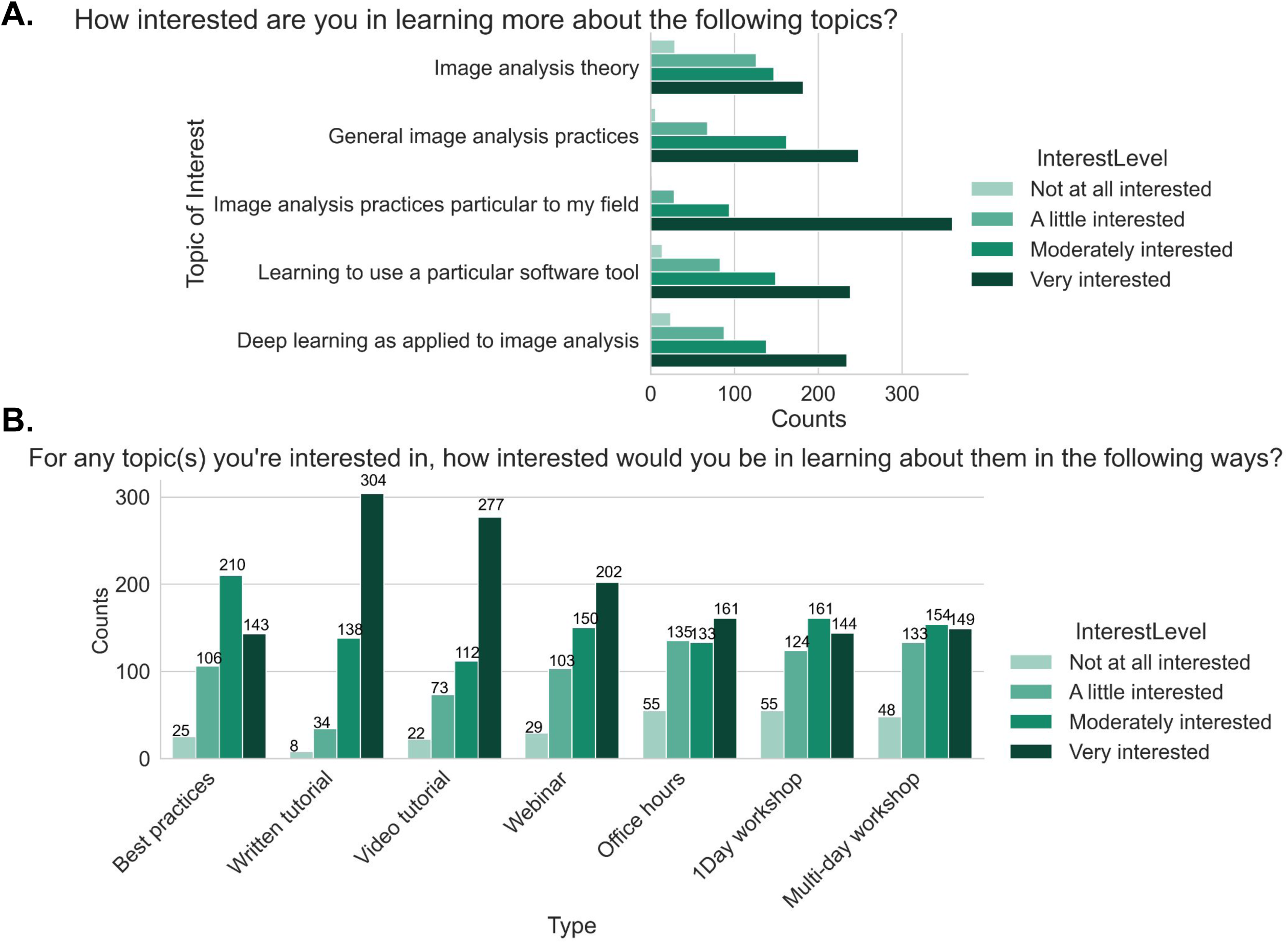
A. Responses to the multiple choice grid question of “How interested are you in learning more about the following topics?”. B. Responses to the multiple choice grid question of “For any topic(s) you’re interested in, how interested would you be in learning about them in the following ways?”.

#### Delivery format of learning material

To help trainers and developers better prioritize particular types of educational material, we asked “*For any topic(s) you’re interested in, how interested would you be in learning about them in the following ways?*”, via a multiple choice grid question. Survey participants in general prefer written step-by-step and video tutorials. The next most preferred methods of learning in order were interactive webinars, best practices articles, and office hours with an expert (Figure 8B). While we hypothesized that we would see differences in preferred format among our “imaging”, “balanced”, and “analyst” groups, we saw fewer differences than we expected between those groups (Supplementary Figure 2A). The results indicated the interest for the written and video tutorial, and interactive webinars are higher between the “imaging” and “balanced” groups at the “moderately” and “very” interested level, while the “analyst” group preferred written tutorial and best practices articles the most. The preferences did not substantially differ between trainees and non-trainees (Supplementary Figure 2B). As we know job role did not perfectly correlate with either existing computational skills or comfort in developing new computational skills, we also wanted to assess how those attributes drove interest in particular types of content; we therefore binned users for each of these categories into “Low” (answers of 1 and 2), “Medium” (answers of 3, 4, and 5), or “High” (answers of 6 and 7). We did not find as much variation in interest level when broken down by comfort level in developing new computational skills (Supplementary Figure 3) as when broken down by computational skills (Supplementary Figure 4); the interest in written material remains high across all demographics but as the level of computational skills increases, the preference for best practices articles increases while the preference for video tutorials or more interactive offerings such as office hours and interactive webinars decreases.

#### Components of well-received learning materials

We asked participants about their past experiences with workshops and conferences. Survey participants have mostly attended workshops, tutorials, and conferences on imaging and image analysis, and they specifically found NEUBIAS (Network of European Bioimage Analysts), Fiji/ImageJ, OME (Open Microscopy Environment), CellProfiler, and Imaris workshops very helpful (Figure 9A). Robert Haase’s workshops on YouTube along with the Analytical and Quantitative Light Microscopy (AQLM) course at Marine Biology Laboratory, and the Quantitative Imaging course at Cold Spring Harbor Laboratory (CSHL) were among the responses as well. When asked what made those particular resources beneficial, respondents highlighted that those workshops provided image analysis basics and theory, step-by-step and hands-on approaches, real-time feedback, and access to experts while developing a workflow.

**Figure 9.**
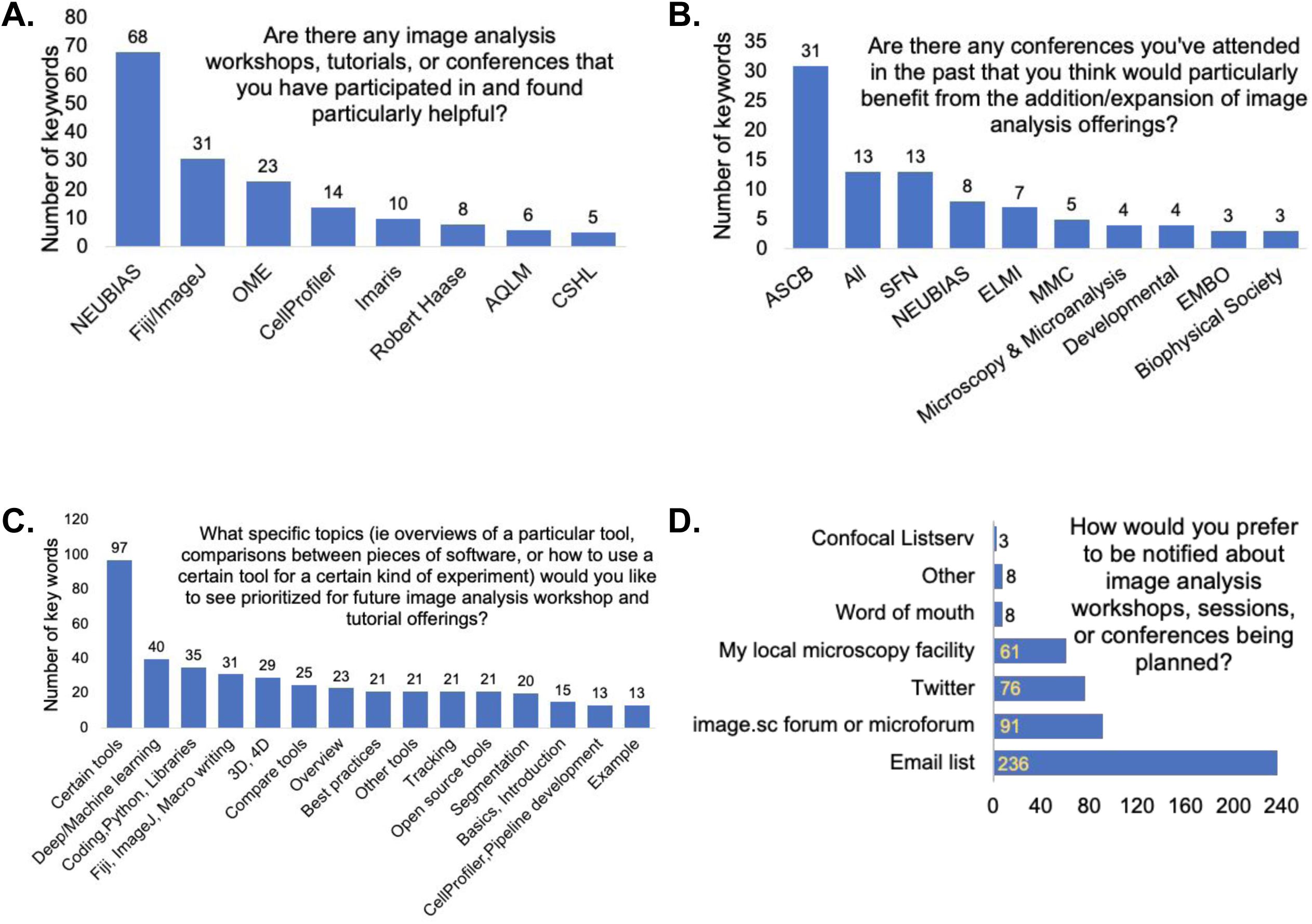
A. Keyword analysis of the open-ended question of “Are there any image analysis workshops, tutorials, or conferences that you have participated in and found particularly helpful? If yes, what made them beneficial?”. B. Keyword analysis of the open-ended question of “Are there any conferences you’ve attended in the past that you think would particularly benefit from the addition/expansion of image analysis offerings?”. C. Keyword analysis of the open-ended question of “What specific topics (i.e. overviews of a particular tool, comparisons between pieces of software, or how to use a certain tool for a certain kind of experiment) would you like to see prioritized for future image analysis workshop and tutorial offerings?”. D. Responses to the multiple choice question of “How would you prefer to be notified about image analysis workshops, sessions, or conferences being planned?”

When asked to name conferences that could benefit from the addition/expansion of image analysis offerings, the most common responses included the following: the American Society for Cell Biology (ASCB), Society for Neuroscience (SFN), Network of European BioImage Analysts (NEUBIAS), European Light Microscopy Initiative (ELMI), and the Microscience Microscopy Congress (MMC) (Figure 9B). Microscopy and Microanalysis, Developmental Biology, and European Molecular Biology Organization (EMBO) scientific conferences were among the responses. We also asked respondents which subjects they would like to see prioritized for the workshops and tutorials. Most - unsurprisingly - wanted materials for the particular tool with which they work, indicating a strong appetite for practical training, but general suggestions included machine and deep learning, coding, Python, different libraries, ImageJ macro writing, overview of different tools, best practices, and different examples (Figure 9C). Finally, the community’s preferred method of notification for the image analysis conferences and workshops are through email lists, forums (such as image.sc and the microforum (forum.microlist.org)), Twitter, and their local microscopy facility (Figure 9D).

In the final “Any other thoughts?” open answer response, the community asked for easy-to-follow (at your own pace) beginner’s guides, better clarification of best practices, a centralized location to find available image analysis tools, various types of example workflows, interoperable platforms to create a workflow combining different tools, and better tools for hyperspectral image analysis.

## Methods

The 2020 BioImage analysis questionnaire was developed using Google Forms and was distributed in the bioimage community around the world using the image.sc forum, microforum, twitter, as well as confocal, imagej, and BioImaging North America (BINA) listservs. Responses were exported via tables, and duplicates were removed; the results were then graphed in Jupyter Notebook (version 6.1.1)^11^ using Python3 ^12^, matplotlib ^13^, numpy ^14^, pandas ^15^, and seaborn ^16^ libraries or Microsoft Excel (2016 and 2021).

To analyze the short and long answer survey questions, the answers to a specific question were parsed with a Python (3.7) script using PyCharm (2019.3) to find the number of words used in the answers, excluding some commonly used words such as: I, that, in, or, of, at. The outcome file was then checked, and depending on the question, the most used and meaningful words were chosen to create a tag list. The responses were parsed again using the tag list, searching for the number of keywords from the tag list. The final results showed the number of keywords (from the tag list) used in the answers. The corresponding figure was then created based on the counts and tags using Microsoft Excel. The tag names used in the figure were sometimes shortened to fit the figure.

## Future needs

In this survey, we strove to include as many participants as possible from as wide a range of biological training backgrounds and computational skill levels as possible. We posted in venues such as FuturePISlack and microscopy listservs; however, the fact that much of the promotion of the survey was via the Scientific Community Image Forum (forum.image.sc) and Twitter accounts related to open-source tools, means it is probable that the group of biologists least exposed to image analysis may also be least represented in this survey. While we have attempted to address this data limitation by breaking down responses to key questions by a number of variables, this limitation points to a larger issue: how can the bioimage analysis community best reach out to the subset of the biology community that is most computationally uncomfortable or unaware and therefore needs user friendly tools the most? We hope that this work, in addition to other recent works describing bioimage analysis surveys ^1^, will also drive community conversation on the most critical questions to quantify and assess the field going forward.

Image analysis is an essential part of microscopy, and in the same way that users learn to optimize their staining, enhance sample preparation, and select appropriate microscope modalities and settings, image analysis method selection, optimization, and building suitable image analysis workflows must be seen as an integral part of the creation of a successful experimental design. Microscopy core facilities are critical for microscopy knowledge at their various institutions; we therefore hope to increase targeted outreach to those core facility staff to improve their access to high quality image analysis resources to then pass onto their users as an essential part of their microscopy training. Since experimental and quantitative biological courses are sometimes the first exposure of trainees to such concepts, we hope to increase our collaborations with instructors to develop the image analysis section of their curricula.

We broadly encourage scientific meetings and conferences to offer image analysis workshops and introduce their various communities to topics such as imaging and image analysis and promote resources such as the microforum (forum.microlist.org) and the Scientific Community Image Forum. While bioimage analysis experts need and benefit from their own internal conferences to share techniques and best practices, especially in the light of users’ preferences to hear more tailored presentations about the kinds of analysis done in their field we hope in the future to be able to partner with both large, broad scientific conferences and small subfield meetings. These partnerships could be used not only to educate attendees, but to promote future attempts to better gauge the biology community’s needs, allowing our bioimage analysis community to hopefully capture a broader perspective. It would also be helpful for funding agencies and journals where tools are published to incentivize creation of “beginner” resources as a standard part of bioimage analysis tool creation, but at the same time tool developers must be given the resources to learn to create such materials. Efforts to “train trainers”, such as The Carpentries (https://carpentries.org/) and Life Science Trainers (https://lifescitrainers.org/), exist and can hopefully be expanded. Finally, we encourage biology graduate and post graduate training organizations to encourage their trainees to think of microscopy images not just as qualitative but as truly quantitative data sources. Their enthusiastic participation in the image analysis community from the beginning of their use of microscopy will encourage improved community engagement.

The Scientific Community Image Forum, which is sponsored by COBA, partners with more than 50 open-source image analysis tools and scientific organizations devoted to biological imaging. It was designed to be a central “go-to” place for the community to ask questions and share updates, news, and ideas. While membership in the forum has grown from approximately 10,000 to 17,000 users in 2 years, fewer than 1/4th of users in our survey identified it as something they “generally” use to solve image analysis problems, indicating the forum can still have an even larger impact among the community. Anecdotal responses from the survey, as well as our experience training users in image analysis, suggest some users find asking questions on the forum intimidating. To ensure the forum lives up to its fullest potential, the community could consider a number of more systematic steps to reduce barriers to participation, such as expanding the existing templates for common questions to make it even easier for users to know which information developers and experts most likely need to answer their questions or creating a stickied “meta-description” post describing all the tools to make choosing a tool easier for new users. Suggestions for improving the forum are always welcome via the various communication routes for COBA via: Twitter (@COBA_NIH), contact forms on our website (https://openbioimageanalysis.org/), and email.

While the broad expansion of open-source image analysis tools has been an incredible benefit to the community at large, such an “embarrassment of riches” can make it overwhelming for a new user to find the most appropriate tool for their image analysis needs. We applaud efforts for cataloguing existing tools such as the BioImage informatics index from NEUBIAS ^17^, and hope that efforts in this space can be expanded. While the image analysis community is already an exceptionally collaborative space, we encourage tool creators to continue to work to make their tools either directly interoperable with popular existing platforms or easily accessed via well-documented Application Programming Interfaces (APIs), as well as to provide other tools with constructive feedback. Together, this will continue to create a rising tide that lifts all boats and gives users easier entry points into the image analysis ecosystem. We encourage forum members to advertise their outreach activities, such as upcoming trainings and links to their previously recorded workshops. We encourage those who are developing bioimage analysis training resources, especially for new users, to reserve a few minutes to introduce users to forum.image.sc, demonstrating how to search for a specific question and find relevant threads, and how to submit an issue or feature improvement request on Github for a specific tool. In addition, event organizers can advertise upcoming conferences and workshops to specifically target their appropriate audience.

There is a clear need for a more centralized online location for the training material related to image analysis tools and resources for best practices. COBA is working toward creating a centralized place for the currently available training resources and best practices guidelines for the community and look forward to collaborating with others looking to do the same.

## Conclusion

Nearly 500 members of the imaging community from around the world participated in the BioImage Analysis survey, conducted in 2020 through the NIH Center for Open BioImage Analysis (COBA). The most common requests from the participants were for general improvement of tool documentation and for access to easy-to-follow tutorials. There is demand for the community to focus on centralizing and publicizing existing educational resources, as well as improving tutorials for the imaging community. Our data on user preferences for particular formats and types of material should help COBA and other developers decide how to structure and prioritize their efforts.

The growth and increased stability of the bioimage analysis community in the last several years is both a triumph and a testament to the many hours that countless members of the community have contributed. With this survey data, COBA hopes to aid the entire community in celebrating its successes and in prioritizing its goals in both the short and longer terms. While individual tools and approaches will no doubt wax and wane in popularity over time, if we share common goals and plans for achieving them, our community will only continue to grow stronger in the years to come.

## Human subject statement

This study was exempt from IRB approval based on Category 2 (Surveys, interviews, educational tests, and public observations). Full exemption was granted because we agreed to de-identify the data, so the subjects’ identity could not be readily obtained.

## Acknowledgements

We thank the biology and microscopy community for providing answers to the survey questions. We also thank Rebecca L. Ledford for her help in opening and publicizing the survey. We thank the members of the Eliceiri, Carpenter-Singh, and Cimini labs for their input on the questionnaire and assessment of needs in the bioimaging community.

## Author contributions

BAC and ETAD created the survey with editorial input from NJ, KWE and AEC. NJ performed analyses with help and supervision from BAC. The manuscript was written by NJ and BAC, with editorial contributions from ETAD, AEC and KWE. All authors have read and approved the manuscript.

## Data availability statement

The survey questions, responses to each individual question (sorted independently for each question to ensure deidentification), and all code used in creating the results are available at https://github.com/ciminilab/2021_Jamali_submitted.

---

**Supplementry Figure 1.**
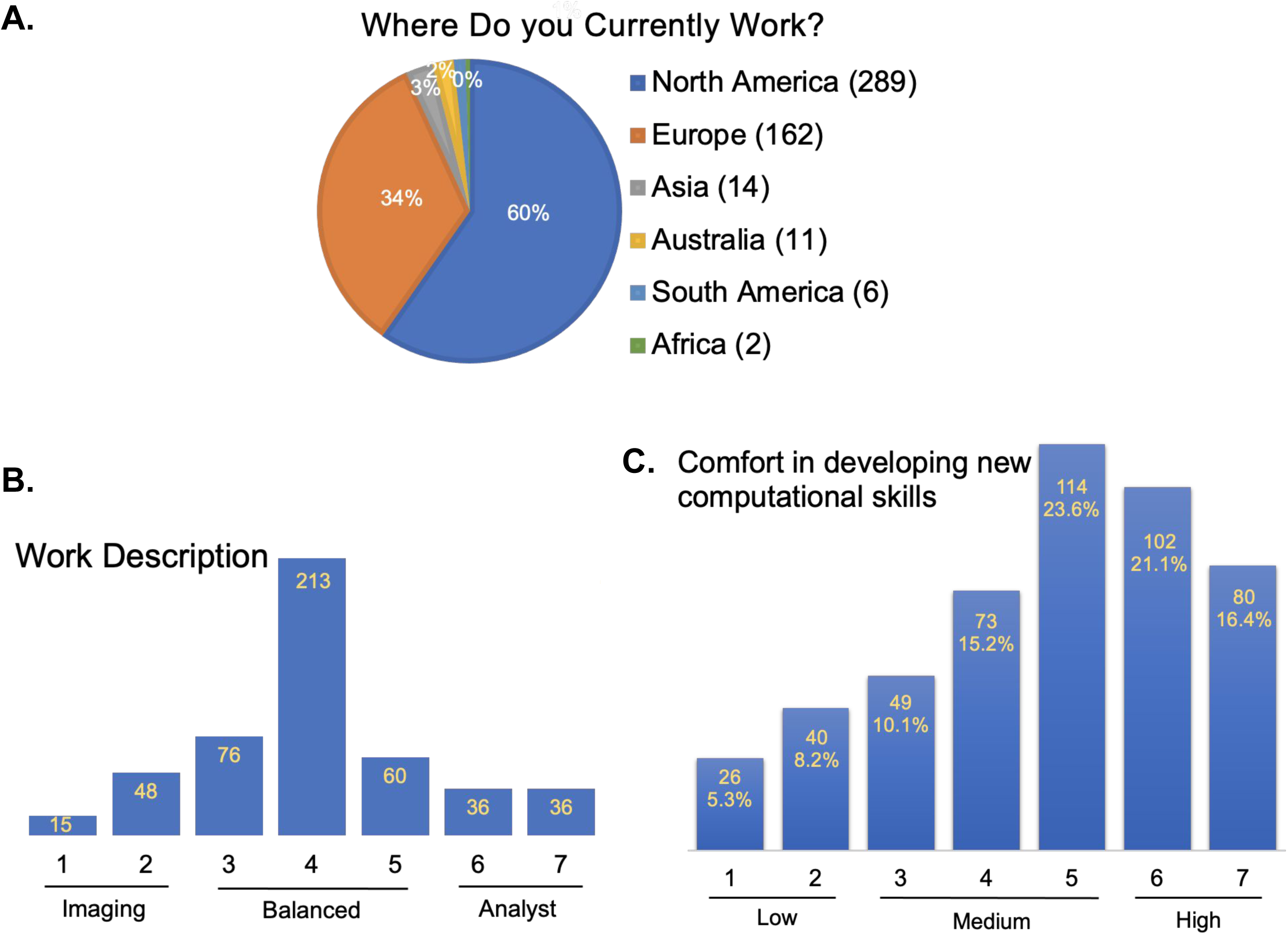
A. Responses to the multiple choice question of “Where do you currently primarily work?” B. Responses to the question of “How would you describe your work?” along a linear scale, with 1 representing *nearly entirely imaging* (sample prep, optimizing/deciding on imaging modalities, acquiring images, etc), and 7 representing *nearly entirely image analysis* (finding the right tools to analyze a particular experiment, optimizing the analysis, data mining). Responses were categorized into three groups: “Imaging” (scores 1&2), “Analyst” (scores 6&7), and “Balanced” (scores 3 to 5). C. Responses to the question of “How would you rate your comfort in developing new computational skills?” along a linear scale, with 1 representing very uncomfortable and 7 representing *very comfortable.* Responses were categorized into three groups of “Low” (scores 1&2), “Medium” (scores 3 to 5), and “High” (scores 6&7).

**Supplementry Figure 2.**
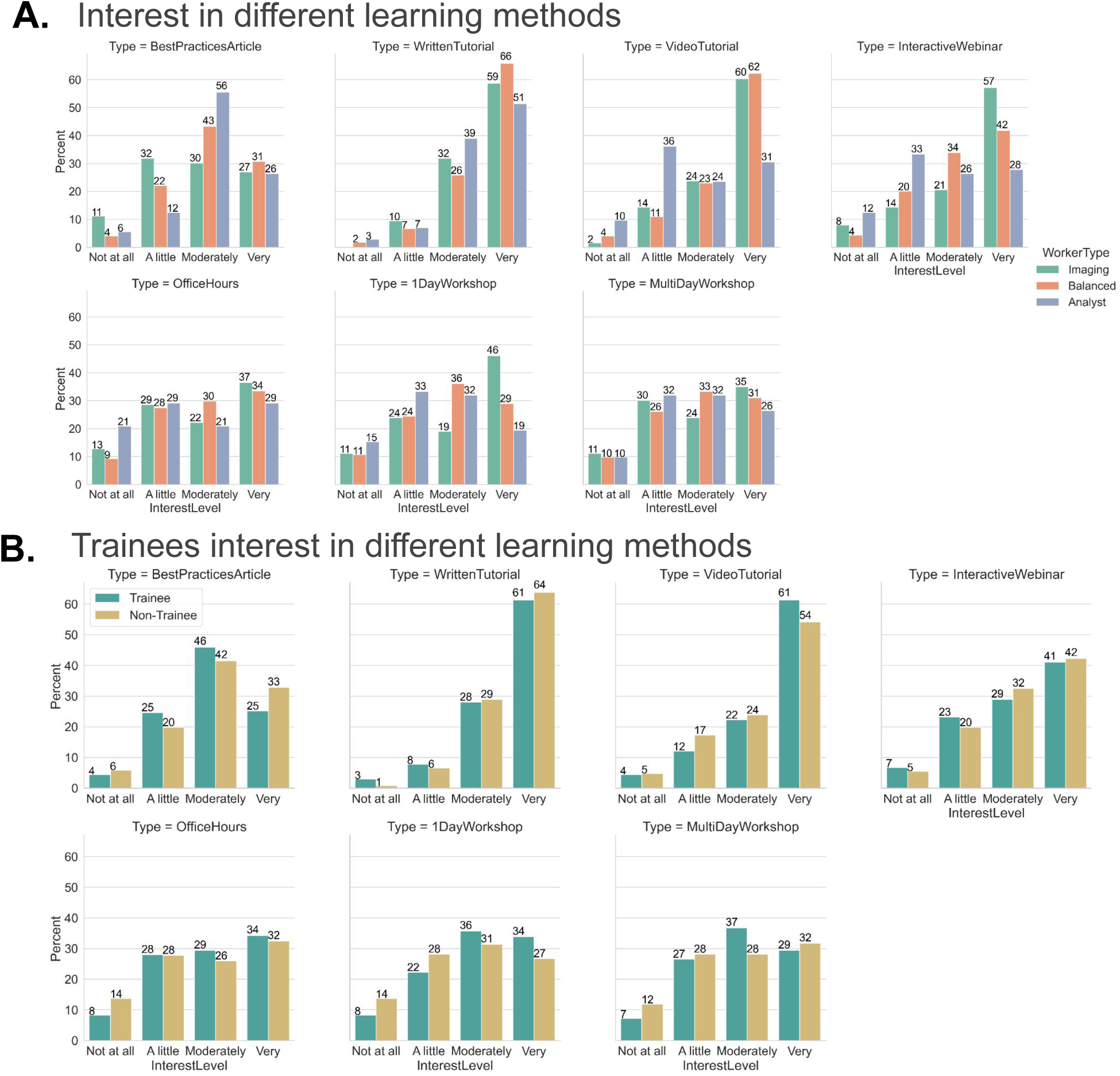
A. Cross-matching of the preferences in learning methods (graphed in Figure 8B) with the work duties (graphed in Supplementary Figure 1B), and separating them based on the level of interest (not at all, a little, moderately, and very). Percentages were calculated independently for each subgroup + method combination. B. Cross-matching of the preferences in learning methods (graphed in Figure 8B) with the trainee group, and separating them based on the level of interest (not at all, a little, moderately, and very). Percentages were calculated independently for each subgroup + method combination.

**Supplementry Figure 3.**
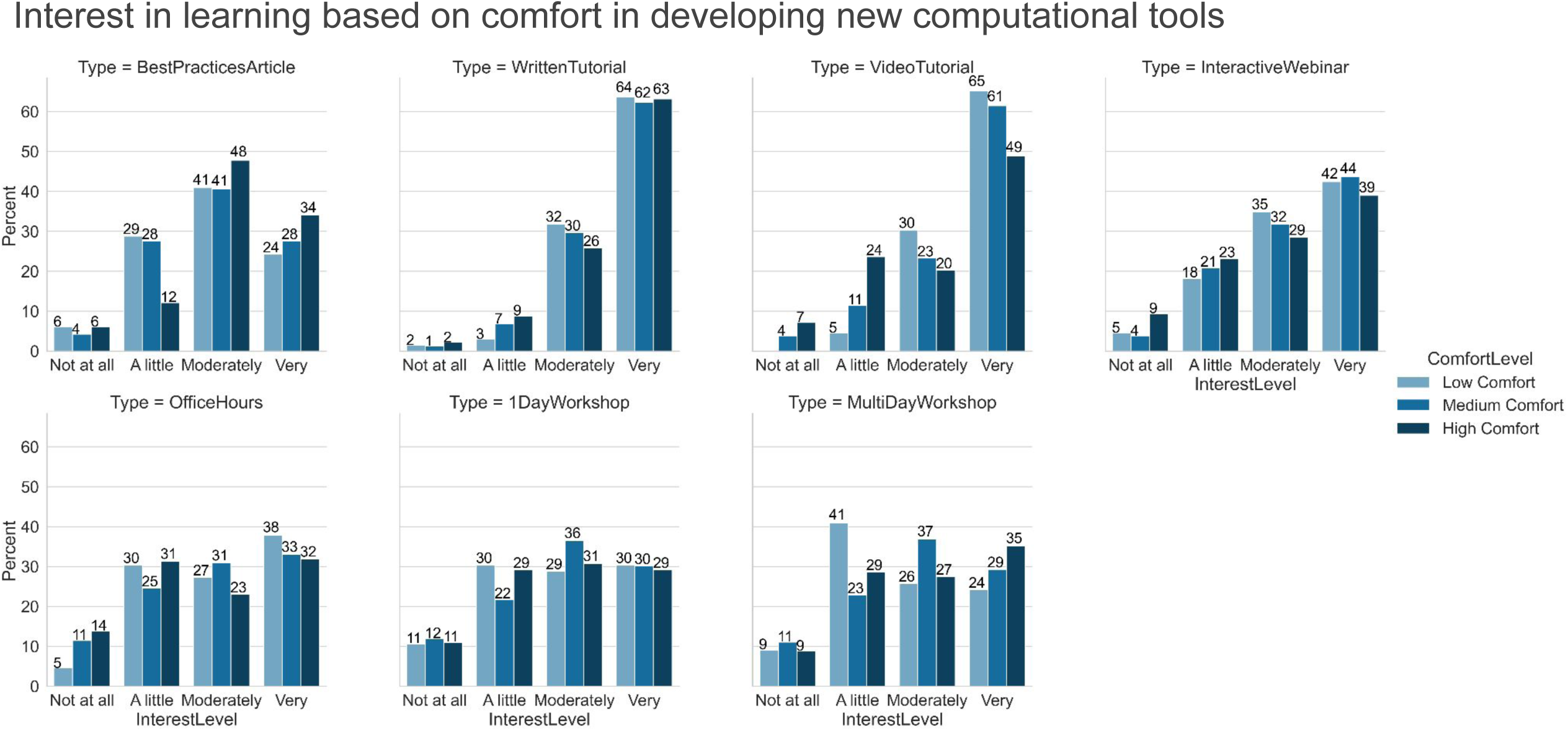
Breakdown of the preferences in learning methods (graphed in Figure 8B) by comfort level in developing new computational skills (graphed in Supplementary Figure 1C). Percentages were calculated independently for each subgroup + method combination.

**Supplementry Figure 4.**
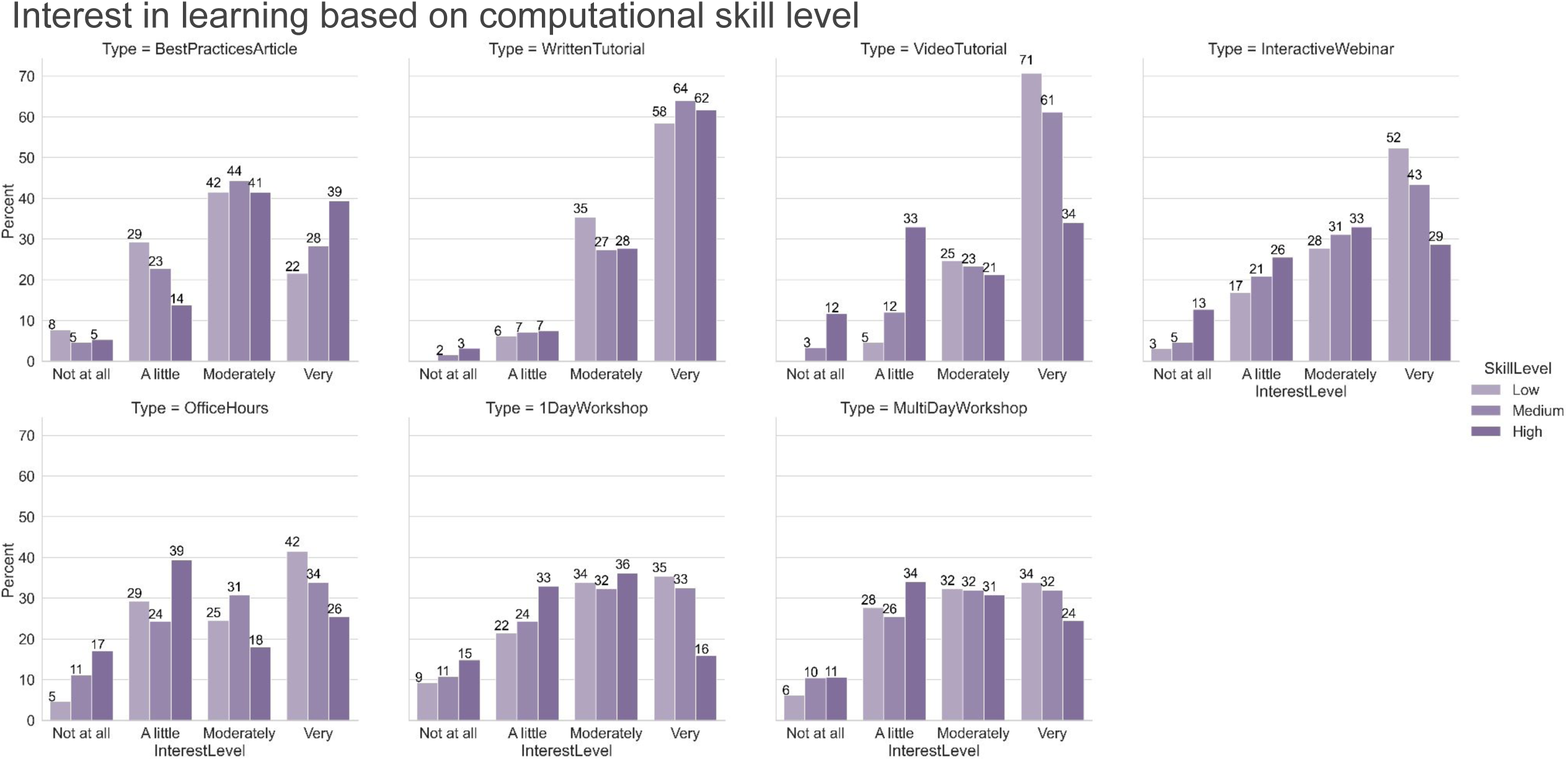
Breakdown of the preferences in learning methods (graphed in Figure 8B) by level of computational skill). Percentages were calculated independently for each subgroup + method combination.

